# Neuropeptide Y signaling regulates recurrent excitation in the auditory midbrain

**DOI:** 10.1101/2023.05.16.540954

**Authors:** Marina A. Silveira, Audrey C. Drotos, Trinity M. Pirrone, Trevor S. Versalle, Amanda Bock, Michael T. Roberts

**Affiliations:** Kresge Hearing Research Institute, Department of Otolaryngology – Head and Neck Surgery, University of Michigan, Ann Arbor, Michigan 48109; Macalester College, St. Paul, Minnesota 55105; Department of Molecular and Integrative Physiology, University of Michigan, Ann Arbor, Michigan, 48109

## Abstract

Neuropeptides play key roles in shaping the organization and function of neuronal circuits. In the inferior colliculus (IC), which is located in the auditory midbrain, Neuropeptide Y (NPY) is expressed by a large class of GABAergic neurons that project locally as well as outside the IC. The IC integrates information from numerous auditory nuclei making the IC an important hub for sound processing. Most neurons in the IC have local axon collaterals, however the organization and function of local circuits in the IC remains largely unknown. We previously found that neurons in the IC can express the NPY Y1 receptor (Y_1_R^+^) and application of the Y_1_R agonist, [Leu^31^, Pro^34^]-NPY (LP-NPY), decreases the excitability of Y_1_R^+^ neurons. To investigate how Y_1_R^+^ neurons and NPY signaling contribute to local IC networks, we used optogenetics to activate Y_1_R^+^ neurons while recording from other neurons in the ipsilateral IC. Here, we show that 78.4% of glutamatergic neurons in the IC express the Y1 receptor, providing extensive opportunities for NPY signaling to regulate excitation in local IC circuits. Additionally, Y_1_R^+^ neuron synapses exhibit modest short-term synaptic plasticity, suggesting that local excitatory circuits maintain their influence over computations during sustained stimuli. We further found that application of LP-NPY decreases recurrent excitation in the IC, suggesting that NPY signaling strongly regulates local circuit function in the auditory midbrain. Together, our data show that excitatory neurons are highly interconnected in the local IC and their influence over local circuits is tightly regulated by NPY signaling.

## Introduction

Local circuits play a critical role in encoding, amplifying and transforming inputs in the brain (Saka et al., 2002; Burke et al., 2017; Geiller et al., 2022). In the auditory midbrain, the inferior colliculus (IC) contains an extensive network of local axons formed by both excitatory and inhibitory neurons (Oliver et al., 1991; Sturm et al., 2014, 2017; Ito et al., 2016). The IC acts as a hub in the central auditory pathway (Adams, 1979; Cant and Benson, 2006, 2007), integrating and transforming many ascending and descending inputs and, in turn, providing major projections to the thalamocortical system and auditory brainstem (Winer et al., 1996; Peruzzi et al., 1997; Goyer et al., 2019; Kreeger et al., 2021; Anair et al., 2022). Although anatomical reports indicate that most neurons in the IC have local axon collaterals (Oliver et al., 1991; Ito et al., 2016), most studies have focused on understanding the long-range projections of IC neurons. Thus, the organization and function of local IC circuits remains poorly understood.

The main barrier towards understanding the function and organization of local IC circuits is that it was not possible to identify and manipulate specific classes of IC neurons (Peruzzi et al., 2000; Palmer et al., 2013; Beebe et al., 2016). However, in the past few years two classes of molecularly distinct glutamatergic neurons have been identified: vasoactive intestinal peptide (VIP (Goyer et al., 2019)) and cholecystokinin (CCK, (Kreeger et al., 2021)). Additionally, we discovered that Neuropeptide Y (NPY) is a molecular marker for a large class of GABAergic principal neurons in the IC (Silveira et al., 2020).

NPY neurons express the peptide NPY, suggesting that this neuropeptide is intrinsically produced and released in the IC. NPY is a powerful neuromodulator known in other brain regions to work in concert with GABA to shape neuronal activity (Colmers and Bleakman, 1994; Bacci et al., 2002; van den Pol, 2012; Gøtzsche and Woldbye, 2016; Li et al., 2017). This suggests the intriguing possibility that NPY signaling regulates auditory processing across spatial and temporal scales not possible with GABAergic signaling alone. Indeed, we previously showed that a subset of glutamatergic neurons in the IC express the NPY Y_1_R receptor (Y_1_R^+^), and application of the Y_1_R agonist [Leu^31^, Pro^34^]-NPY (LP-NPY) led to hyperpolarization of Y_1_R^+^ IC neurons (Silveira et al., 2020). Y_1_R^+^ neurons are thought to be glutamatergic neurons since they do not express the GABA-synthetic enzyme GAD67, and they likely encompass more than one class of glutamatergic neurons since they exhibit heterogenous intrinsic physiology (Silveira et al., 2020). The presence of Y_1_R^+^ neurons in the IC raises the hypothesis that NPY is a modulator of local network activity, however how Y_1_R^+^ neurons and NPY signaling shape local circuit function in the IC is unknown.

Here, we used whole-cell patch clamp recordings and optogenetics in brain slices from Y_1_R-Cre x Ai14 mice, in which Y_1_R^+^ neurons are identified by tdTomato expression, to test the hypothesis that NPY signaling modulates local circuits in the IC. Our data show that the 78.4% of glutamatergic neurons in the IC express *Npy1r* mRNA and that Y_1_R^+^ neurons can also express *VIP* and/or *CCK*. We also show that Y_1_R^+^ neurons are densely interconnected in local IC circuits, providing extensive excitatory inputs to other neurons in the IC.

Contrary to what has been shown for ascending inputs to the IC (Wu et al., 2004), local Y_1_R^+^ synapses exhibit little short-term synaptic plasticity suggesting a balance between short-term synaptic depression and short-term synaptic facilitation. We further found that application of LP-NPY decreased recurrent excitation in the IC. Together, our results reveal that Y_1_R^+^ neurons in the IC form highly interconnected local circuits and that the excitability of these circuits is regulated by NPY signaling. Thus, we propose that NPY signaling is a major modulator of the auditory computations performed by local circuits in the auditory midbrain.

## Materials and Methods

### Animals

The experiments performed here were approved by the University of Michigan Institutional Animal Care and Use Committee and were in accordance with National Institutes of Health’s *Guide for the Care and Use of Laboratory Animals*. Mice had ad libitum access to food and water and were maintained on a 12 h day/ night cycle. Npy1r^cre^ mice (B6.Cg-Npy1r^tm1.1(cre/GFP)Rpa^, The Jackson Laboratory, stock # 030544 (Padilla et al., 2016)), were crossed with Ai14 Cre-reporter mice (B6.Cg-Gt(ROSA)26Sortm14(−CAG-tdTomato)Hze; The Jackson Laboratory, stock # 007914 (Madisen et al., 2010)) to allow identification of Y_1_R^+^ neurons by tdTomato fluorescence (Y_1_R-Cre x Ai14). Both mouse lines were on a C57BL/6J background. Since C57BL/6J mice are subject to age-related hearing loss due to the *Cdh23*^*ahl*^ mutation, experiments were performed in mice aged P24 -82 to avoid potential effects of age-related hearing loss (Noben-Trauth et al., 2003). Mice of either sex were used for all experiments.

### RNAScope in situ hybridization assay and analysis

Two *in situ* hybridization assays were performed. For both assays, we used the RNAScope Multiplex Fluorescence V2 kit (Advanced Cell Diagnostics, catalog # 320850, (Wang et al., 2012)). Brain slices were prepared using the *fresh-frozen* method. Mice were deeply anesthetized with isoflurane, and brains from three mice (two males and one female Y_1_R-Cre x Ai14 aged P59 for assay #1 and two females and one male C57BL/6J aged P55 for assay #2) were quickly dissected, frozen in dry ice, and maintained at -80 °C until it was time to slice. Prior to slicing, brains were equilibrated at -20 °C for one hour and 15 μm sections were collected using a cryostat and mounted on Superfrost Plus slides (Fisher Scientific, catalog # 22037246). RNAScope assays were performed following the recommendations of the manufacturer. In brief, slices were fixed in 10% neutral buffered formalin (Sigma-Aldrich, catalog # HT501128), dehydrated in increasing concentrations of ethanol, followed by the drawing of a hydrophobic barrier around the sections. Next, hydrogen peroxide was used to block endogenous peroxidase. For hybridization, we used the following combinations of probes: (1) *tdTomato, Vglut2* and *Npy1r* and, (2) *VIP, CCK* and *Npy1r*. All probes, positive controls and negative controls were incubated for 2 hours followed by the amplification (AMP 1 – 3) step. After signal was developed using the appropriate HRP, Opal dyes (1:1000) were assigned for each channel: Assay # 1: *tdTomato* expression was identified by Opal 690 (Akoya Bioscience, catalog # FP1497001KT), *Vglut2* expression was identified by Opal 570 (Akoya Bioscience, catalog # FP1488001KT) and *Npy1r* expression was identified by Opal 520 (Akoya Bioscience, catalog # FP1487001KT); assay # 2: *VIP* expression was identified by Opal 520 (Akoya Bioscience, catalog # FP1487001KT), *CCK* expression was identified by Opal 570 (Akoya Bioscience, catalog # FP1488001KT) and *Npy1r* expression was identified by Opal 690 (Akoya Bioscience, catalog # FP1497001KT). Following staining with DAPI, slices were coverslipped using ProLong Gold antifade mountant (Fisher Scientific, catalog # P36934). Images were collected within two weeks after the assay. Representative sections (including caudal, mid-rostrocaudal, and rostral, 3 sections per mouse) were imaged using a 0.75 NA 20X objective at 2 μm depth intervals on a Leica TCS SP8 laser scanning confocal microscope (Leica Microsystems). For the validation of the mouse line (assay # 1), quantification was performed manually using FIJI (Image J, National Institutes of Health (Schindelin et al., 2012)). A grid was randomly placed over every image and quantification was performed every 4^th^ – 6^th^ grid square using the multi-point tool by placing a marker on the top of each cell. Channels of each color were quantified separately to avoid bias. To test whether *Npy1r* expressing neurons express *VIP* and/or *CCK* (assay # 2) we looked for representative examples of cells expressing a combination of the three different mRNAs.

### Intracranial Virus Injection

To investigate the function of IC local circuits, 17 Y_1_R-Cre x Ai14 mice (9 males and 8 females) were injected in the right IC with the recombinant adeno-associated virus (rAAV) rAAV1/ Syn-Flex-Chronos-GFP (Lot # AV6551B, UNC Vector Core, Addgene # 62725, titer 2.8 × 10^12^ vg/ml (Klapoetke et al., 2014)). For short-term synaptic plasticity experiments and recurrent excitation experiments, 21 Y_1_R-Cre x Ai14 mice (14 males and 7 females) were injected in the right IC with either rAAV1/ Syn-Flex-Chronos-GFP or rAAV9-CAG-DIO-ChroME-ST-P2A-H2B-mRuby3 (Addgene # 108912, titer ranging from 2.7 × 10^12^ to 2.7 × 10^13^ gc/ml (Mahn et al., 2018)). Surgeries were performed using standard aseptic techniques in males and females, aged P25 – P68 as previously described (Goyer et al., 2019; Silveira et al., 2020). In brief, mice were deeply anesthetized using 3% isoflurane and placed in a stereotaxic base with a homeothermic heating pad to maintain body temperature. For the remainder of the surgery, isoflurane was dropped to 1 – 2 % and breathing pattern was monitored. Carprofen (5 mg/kg, CarproJect, Henry Schein Animal Health) was injected subcutaneously. After exposing the skull with a rostrocaudal incision, a unilateral craniotomy was performed in the right IC using a micromotor drill (K.1050, Foredom Electric) with a 0.5 mm burr (Fine Science Tools). Injections were made in two penetrations using coordinates were defined relative to lambda and relative to the surface of the skull. Injection # 1: 900 mm caudal, 1000 mm lateral and 1800 mm deep and injection # 2: 900 mm causal, 1200 mm lateral and 1800 mm deep. Glass capillary used for injections (catalog # 3-000-203-G/X, Drummond Scientific) were pulled with a P-1000 microelectrode puller (Sutter Instrument). The injection tip was back filled with mineral oil and front filled with the virus of interest. The amount of virus injected ranged from 50 – 300 nl, however most mice were injected with 100 nl of virus. At the end of the procedure, the scalp was sutured using Ethilon 6-0 (0.7 metric) nylon sutures (Ethicon). The analgesic lidocaine hydrochloride (2%, 0.5ml, Akorn) was applied to the incision. Mice were monitored for 10 days. Electrophysiological recordings were performed 2 -4 weeks after the injection.

### Brain slice preparation

To characterize the intrinsic physiology of Y_1_R^+^ neurons and to evaluate local projections mediated by Y_1_R^+^ neurons, we performed whole-cell patch clamp recordings targeted to either Y_1_R^+^ or Y_1_R^-^ neurons that were identified in Y_1_R-Cre x Ai14 mice by the presence or lack of tdTomato expression. To prepare brain slices, mice were decapitated after being deeply anesthetized with isoflurane. The brain was carefully removed, and the IC was dissected in 34 °C ACSF containing (in mM): 125 NaCl, 12.5 glucose, 25 NaHCO_3_, 3 KCl, 1.25 NaH_2_PO_4_, 1.5 CaCl_2_, 1 MgSO_4_, 3 sodium pyruvate, and 0.40 L-ascorbic acid. ACSF was bubbled to a pH of 7.4 with 5 % CO_2_ in 95 % O_2_ for at least 30 min prior to dissection. Coronal IC slices (200 μm) were prepared using a vibrating microtome (VT1200S, Leica Biosystems). Slices were incubated at 34 °C for 30 min in ACSF bubbled with 5 % CO_2_ in 95 % O_2_. Unless otherwise noted, all reagents were obtained from Thermo Fisher Scientific.

### Electrophysiological recordings

For whole-cell current clamp and voltage clamp recordings, slices were transferred to a bath chamber that was continuously perfused with 34 – 35 °C ACSF bubbled with 5 % CO_2_ in 95 % O_2_ at 2 ml/min. Y_1_R^+^ and Y_1_R^-^ neurons were identified with epifluorescence using a Nikon FN1 microscope or an Olympus BX51WI microscope. Recording pipettes were pulled using a P-1000 microelectrode puller (Sutter Instrument) from borosilicate glass capillaries (outer diameter 1.5 mm, inner diameter 0.86 mm, catalog # BF150-86-10, Sutter Instrument). Internal solution to fill the pipettes contained (in mM): 115 K-gluconate, 7.73 KCl, 0.5 EGTA, 10 HEPES, 10 Na_2_-phosphocreatine, 4 MgATP, 0.3 NaGTP and 0.1% biocytin (w/v). The pH was adjusted to 7.3 with KOH and osmolality to 290 mmol/kg with sucrose. Pipettes with resistances from 2.5 – 5.5 MΩ were used for recordings.

Whole-cell current clamp recordings were performed using a BVC-700A amplifier (Dagan). Data were acquired using custom software written in IgorPro, low-pass filtered at 10 kHz and sampled at 50 kHz with a National Instruments PCIe-6343 data acquisition board. Pipette capacitance and series resistance were corrected using the bridge balance circuitry of the BVC-700A amplifier. Recordings with series resistance >20 MΩ and recordings with series resistance that changed more than 15 % were not included in analysis. To assess input resistance a series of 100 ms current steps that hyperpolarized the membrane potential from just below rest to -100 mV was applied. The peak and steady-state voltage responses were calculated for each step. A voltage versus current plot was generated and the peak and steady-state input resistances were calculated based on the slope of a linear fit. To calculate the membrane time constant fifty, 100 -300 ms current steps that resulted in 2 -6 mV membrane hyperpolarization were applied. Membrane time constant was obtained by fitting an exponential function to each response and obtaining the median τ. Liquid junction potential was -11 mV and was corrected in all cases.

Whole-cell voltage clamp recordings were made using an Axopatch 200A amplifier. Series resistance compensation was performed using 70 - 80% prediction and correction. Recordings with series resistance >20 MΩ or that changed more than 15 % during the recording were not included in the final analysis. Internal solution was the same one used for current clamp experiments and recordings were performed at -81 mV with the liquid junction potential of -11 mV corrected in all cases.

### Channelrhodopsin-assisted circuit mapping (CRACM)

After injections, we allowed 2 – 4 weeks for Chronos or ChroME expression. Brain slices were prepared, and recordings were performed under red light to limit Chronos or ChroME activation. Recordings were targeted to Y_1_R^+^ or Y_1_R^-^ neurons and the identification of neurons was done as fast as possible to avoid activation of optogenetic opsins. Chronos or ChroME were activated by brief pulses of 470 nm light emitted by a blue LED coupled to the epifluorescence path of the microscope. Light was delivered to the brain slice through a 0.80 NA 40x water immersion objective. Optical power densities ranged from 6 to 48 mW/mm^2^. Recording sweeps with light presentations were repeated 20 to 100 times. Sweeps were collected at 20 second intervals using a light duration of 0.5 -5 ms, but most of the data was acquired using 1 – 2 ms light pulse. Sweeps that did not exhibit a light evoked EPSP were excluded from analysis. A minimum of 13 sweeps with an EPSP was required to include a cell in the analysis. To determine the receptors contributing to excitatory postsynaptic potentials (EPSPs), 10 μM NBQX disodium salt (AMPA receptor antagonist, Hello Bio, Hello Bio Cat#: HB0442) and 50 μM D-AP5 (NMDA receptor antagonist, Hello Bio Cat#: HB0225) were used. Drugs were bath applied for at least 10 minutes before data were collected.

### Short-term synaptic plasticity

For short-term synaptic plasticity experiments, 5 μM gabazine (also called SR95531 hydrobromide, GABA_A_ receptor antagonist, Hello Bio, Cat #: HB0901) and 1 μM strychnine hydrochloride (Sigma-Aldrich, Cat #: S8753) were added to the bath to block feedforward inhibition. A total of five EPSPs were elicited using 20 Hz trains of 1 – 2 ms light pulses. For analysis, cursors were placed at the beginning and end of the EPSP to determine the peak amplitude of the first EPSP. The peak amplitude of the following EPSPs was determined using the foot-to-peak method in which the peak amplitude was measured relative to the most negative point attained following the peak of the preceding EPSP. Paired pulse ratios (PPR) were defined by calculating the ratio of (average amplitude of EPSP_n_)/(average amplitude of EPSP_1_) (Kim and Alger, 2001). Sweeps that exhibited recurrent excitation were excluded from the analysis.

### Recurrent excitation recordings

To enhance recurrent excitation in brain slices, 5 μM gabazine and 1 μM strychnine were included in the ACSF solution (Tu et al., 2005). Activation of excitatory inputs mediated by Y_1_R^+^ neurons was achieved by transfecting Y_1_R^+^ neurons with Chronos or ChroME and using brief flashes of blue light to activate Y_1_R^+^ inputs. After light presentation (0.5 – 5 ms), a large recurrent excitation was elicited causing the membrane to reach threshold and fire action potentials. After recording at least 20 sweeps in control condition, LP-NPY (500 nM diluted in 20 ml of ACSF, Tocris Cat #: 1176) was bath applied for ∼10 min. After application of LP-NPY, the bath solution was returned to control ACSF and washout responses were measured 10 – 30 min later. The number of action potentials was calculated in the control condition, during application of LP-NPY and washout. Action potentials were counted using a custom threshold-crossing algorithm in Igor Pro 9 (Wavemetrics). Threshold was defined by eye for each cell. To determine the area under the curve, data were median filtered using a 4000 sample (80 ms) smoothing window to remove action potentials while retaining the underlying waveform of the depolarization.

To determine the effect of LP-NPY on recurrent excitatory currents in the local IC, experiments were performed in voltage clamp and brief pulses of blue light were presented to elicit a long-lasting excitatory current. Only cells with a depolarizing current that lasted more than 40 ms were included in the analysis. The area under the curve was calculated for control, LP-NPY and washout. To test if responses were synaptically mediated, 10 μM NBQX and 50 μM D-AP5 were bath applied for at least 10 minutes and additional responses were collected.

To determine the effect of LP-NPY on recurrent excitation, values of action potential number and the area under the curve were plotted in IgorPro 9 (Wavemetrics). A custom algorithm was used to identify the most negative value during LP-NPY application. We than calculated the average across 9 sweeps during baseline, LP-NPY application and washout. Baseline and washout were defined relative to the most negative peak of LP-NPY effect. Baseline was calculated using the average value of action potentials or area under the curve across 760 – 920 seconds (12.6 – 15.3 minutes) prior to the LP-NPY peak effect. Washout was calculated 540 – 700 seconds (9 – 11.6 minutes) after the LP-NPY peak effect (illustrated as gray bars on **Figures 8D-F** and **9E**).

### Statistics

Statistical analyses were performed in Igor Pro 9 (Wavemetrics), R 4.1.0 (The R Project for Statistical Computing) and MATLAB R2021a (MathWorks). Data were analyzed using the estimation statistics approach (Bernard, 2019; Calin-Jageman and Cumming, 2019; Calin-Jageman, 2022), which has been previously detailed in a study from our lab (Rivera-Perez et al., 2021). For the comparison of the intrinsic physiology between adapting and sustained Y_1_R^+^ neurons (**Figure 3**) we used the “independence test,” which is a non-parametric permutation test (Strasser and Weber, 1999). The significance level (α) was adjusted to account for the multiple comparisons using Bonferroni correction. For linear mixed models (LMMs) analysis we used the “lme4” and “lmerTest” packages in R (Bates et al., 2015). For most experiments, effects were considered significant when p < 0.05. Because the baseline area under the curve for the voltage clamp and current clamp experiments, as well as the action potential number, varied largely across cells, we calculated the log of the raw data before running LMM (**Figures 8** and **9**). Principal component analysis (PCA) was done in MATLAB using the *“coeff”* function. The k-means test was performed after PCA using the *“kmeans”* function in MATLAB. The optimal value of k (clusters) was defined as 2 using the “elbow method” (see insert graph in **Figure 3**). The statistical test used for each experiment is described in results.

## Results

### 78.4% of glutamatergic neurons in the IC express Npy1r mRNA

To identify neurons that express the Y_1_R, we used Y_1_R-Cre x Ai14 mice to label Y_1_R^+^ neurons with the tdTomato fluorescent protein (Padilla et al., 2016). To validate that these mice selectively expressed tdTomato in Y_1_R^+^ neurons, we performed *in situ* hybridization using RNAScope with probes targeted to *Npy1r, Vglut2*, and *tdTomato*. Two males and one female mouse aged P59 were used for the assay. Using design-based stereology, we found that 79.8% (3726 out of 4668) of t*dTomato*^*+*^ neurons co-labeled with *Npy1r* (**Figure 1, Table 1**).

**Table 1.** tdTomato^+^ (tdT^+^) neurons are glutamatergic and express Y_1_R.

| Mouse | Slice # | *tdT<sup>+</sup>* | *tdT<sup>+</sup>*<br>*Vglut2<sup>+</sup>* | % *tdT<sup>+</sup>Vglut2<sup>+</sup>*<br>/*tdT<sup>+</sup>* | *tdT<sup>+</sup>*<br>*Npy1r<sup>+</sup>* | % *tdT<sup>+</sup>Npy1r<sup>+</sup>*<br>/*tdT<sup>+</sup>* |
| --- | --- | --- | --- | --- | --- | --- |
| Fem P59 | Caudal | 278 | 254 | 91.4 | 180 | 64.7 |
|  | Medial | 371 | 334 | 90.0 | 264 | 71.2 |
|  | Rostral | 432 | 403 | 93.3 | 344 | 79.6 |
| Male P59 | Caudal | 464 | 426 | 91.8 | 331 | 71.3 |
|  | Medial | 421 | 390 | 92.6 | 319 | 75.8 |
|  | Rostral | 260 | 235 | 90.4 | 195 | 75.0 |
| Male P59 | Caudal | 522 | 478 | 91.6 | 444 | 85.1 |
|  | Medial | 1120 | 1026 | 91.6 | 964 | 86.1 |
|  | Rostral | 800 | 748 | 93.5 | 685 | 85.6 |
| Total |  | 4668 | 4294 | 92.0 | 3726 | 79.8 |

**Figure 1.**
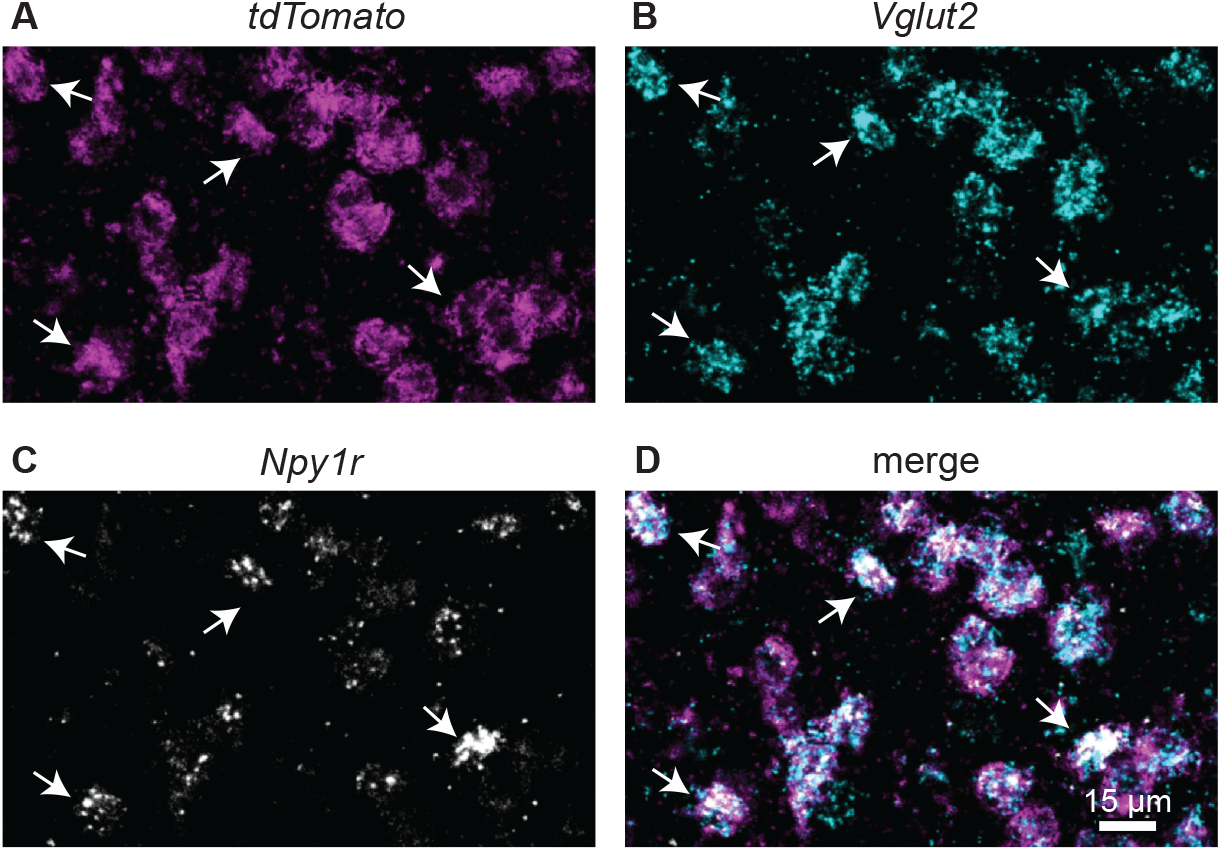
*tdTomato*^*+*^ neurons express *Vglut2* and *Npy1r* mRNA. High magnification confocal images of a coronal IC section from a Y_1_R-Cre x Ai14 mouse showing that *tdTomato*^*+*^ neurons (**A**, magenta) co-label with *Vglut2* (**B**, cyan) and *Npy1r* (**C**, white; merge in **D**). White arrows show examples of neurons that co-label. Scale bar applies to all images.

In our previous study, we showed that only 1.1% of Y_1_R^+^ neurons co-labeled with GAD67, a marker for GABAergic neurons, suggesting that most Y_1_R^+^ neurons are glutamatergic (Silveira et al., 2020). To confirm the neurotransmitter content of Y_1_R^+^ neurons, we used a probe targeted to *Vglut2*, which is a marker of glutamatergic cells in the IC (Ito et al., 2011). We found that 92% (4294 out of 4668) of *tdTomato*^*+*^ neurons express *Vglut2*. In very rare cases (0.4%), cells expressed *Npy1r* without the expression of *tdTomato* (15 out of 3741 *Npy1r* expressing neurons, **Table 2**). Despite the expression of *td-Tomato*, 97.5% of *Npy1r* expressing neurons expressed *Vglut2* (3647 out of 3741). We then quantified the remaining glutamatergic cells (357 cells that did not express *tdTomato* and/or *Npy1r*) and found that *Npy1r* expressing neurons represented 78.4% of glutamatergic cells in the IC (3647 out of 4651, **Table 2**). This was a striking result since it shows that most IC glutamatergic neurons express *Npy1r*, suggesting that NPY signaling plays a major role in regulating excitatory neurons in the IC.

**Table 2.**
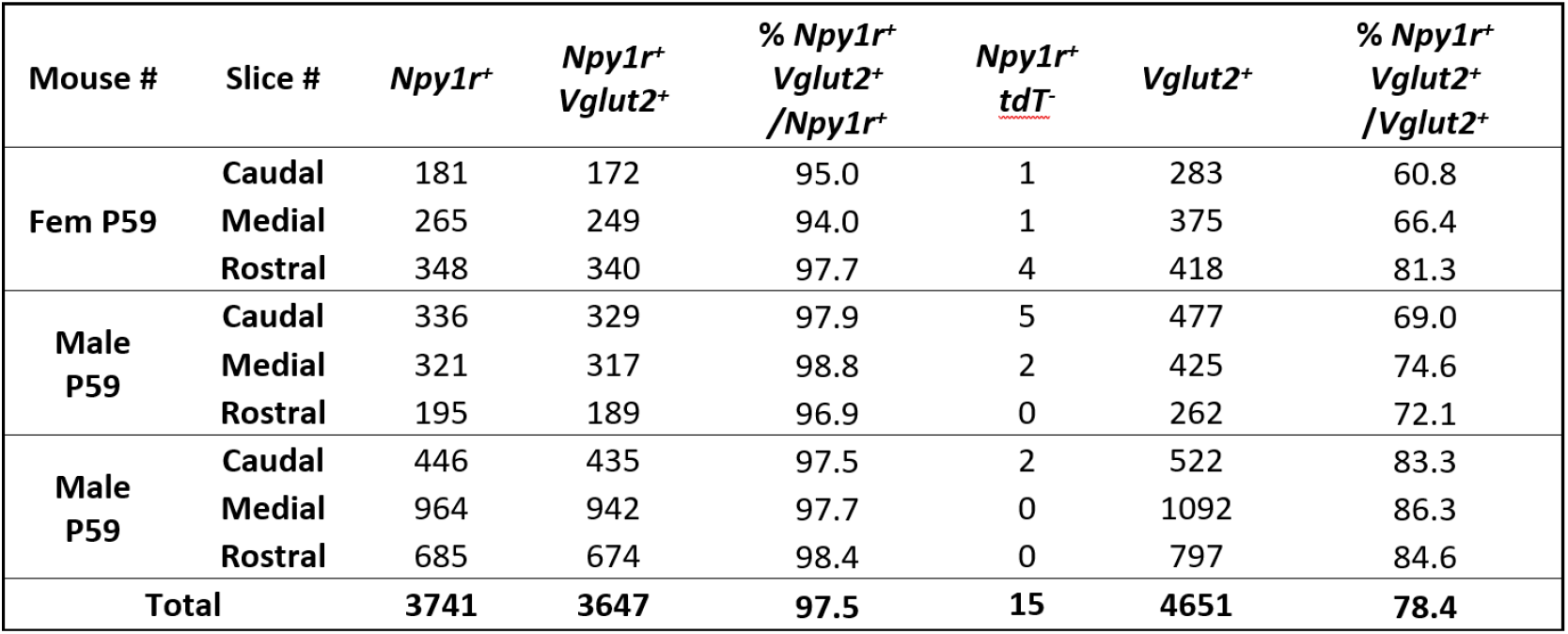
Y_1_R^+^ neurons represent 78.4% of IC glutamatergic cells.

Previous studies described two subclasses of glutamatergic neurons in the IC, identified by the expression of VIP (Goyer et al., 2019) and CCK (Kreeger et al., 2021). To test whether Y_1_R^+^ neurons express *VIP* and/or *CCK*, we next performed *in situ* hybridization with probes targeted to *Npy1r, VIP* and *CCK*. We found that many, but not all, *CCK* and *VIP* neurons expressed *Npy1r*. Interestingly, we also found that *VIP* and *CCK* expression could overlap, with many *VIP* neurons expressing *CCK*, but *VIP*-expressing neurons representing only a subset of *CCK* neurons (**Figure 2**, blue arrow indicates *VIP*^*+*^*CCK*^*+*^*Y*_*1*_*R*^*-*^; white arrow indicates *VIP*^*-*^*CCK*^*+*^*Y*_*1*_*R*^*+*^; magenta arrow indicates *VIP*^*+*^*CCK*^*+*^*Y*_*1*_*R*^*+*^).

**Figure 2.**
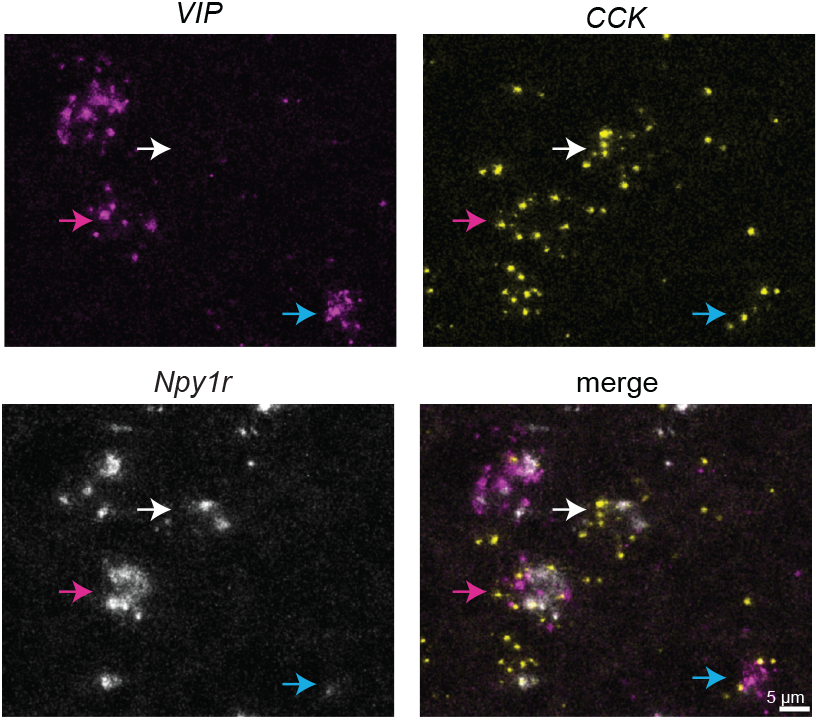
Y_1_R^+^ neurons can express *VIP* and/or *CCK*. High magnification confocal images of a coronal IC section showing *VIP* (magenta), *CCK* (yellow), and *Npy1r* (white) expression. Arrows indicate examples of cells that are *VIP*^*+*^*CCK*^*+*^*Y*_*1*_*R*^*-*^(blue arrow); *VIP*^*-*^*CCK*^*+*^*Y*_*1*_*R*^*+*^(white arrows); and *VIP*^*+*^*CCK*^*+*^*Y*_*1*_*R*^*+*^ (magenta arrow). Scale bar applies to all images.

### Y_1_R^+^ neurons exhibit heterogenous intrinsic physiology

Neurons in the IC exhibit diverse firing patterns, being broadly classified as sustained or adapting neurons (Peruzzi et al., 2000; Goyer et al., 2019; Silveira et al., 2020). By targeting whole-cell current clamp recordings to Y_1_R^+^ neurons in acute brain slices from Y_1_R-Cre x Ai14 mice, we previously showed that Y_1_R^+^ neurons exhibit either sustained or adapting firing patterns in response to depolarizing current steps (Silveira et al., 2020). Here we expanded the recordings (n = 38 cells previously reported in Silveira et al., 2020 plus an additional n = 55 cells from this study). We found that Y_1_R^+^ neurons with sustained or adapting firing patterns exhibited heterogeneous intrinsic properties. Most neurons had a sustained firing pattern in response to a depolarizing current step (60 out of 93 neurons had a spike frequency adaption (SFA) ratio < 2, and 33 out of 93 neurons had an SFA ratio > 2; **Figure 3A,B**; SFA ratio with 5 spikes comparing sustained vs adapting neurons: unpaired independence test, Z = 6.30, *p* < 1e-04; Bonferroni-corrected α = 0.0083. Sustained Y_1_R^+^ neurons had a slower membrane time constant (15.39 ± 4.90 ms) compared to adapting neurons (8.40 ± 4.90 ms, independence test, Z = -4.44, *p* < 1e-04; Bonferroni-corrected α = 0.0083; **Figure 3C**). In response to hyperpolarizing current steps, adapting Y_1_R^+^ had a more prominent sag ratio suggesting a higher expression of HCN channels (steady state/peak, measured from current steps that elicited peak hyperpolarization of -91.0 ± 0.9 mV; voltage sag ratio of 0.74 ± 0.19 for adapting neurons and 0.88 ± 0.14 for sustained neurons, independence test, Z = -3.56, *p* < 1e-04; Bonferroni-corrected α = 0.0083; **Figure 3D**). Rheobase values were lower for sustained neurons (56.18 ± 37.63 pA) compared to adapting neurons (77.94 ± 47.23 pA, independence test, Z = 2.37, *p* = 0.01; Bonferroni-corrected α = 0.0083; **Figure 3E**). Resting membrane potential was similar between adapting and sustained neurons (−67.9 ± 6.5 mV for adapting neurons and -69.3 ± 6.6 mV for sustained neurons, independence test, Z = 0.96, *p* = 0.33; Bonferroni-corrected α = 0.0083; **Figure 3F**). Finally, sustained neurons exhibited a higher input resistance (266.8 ± 99.5 MΩ, measured at the peak of the hyperpolarizing response) compared to adapting neurons (167.9 ± 74.7 MΩ, independence test, Z = -10.57, *p* < 1e-04; Bonferroni-corrected α = 0.0083; **Figure 3G**).

We next performed principal components analysis (PCA) to test whether Y_1_R^+^ neurons can be divided into multiple cell types based on intrinsic physiology. The PCA was performed using the following parameters: membrane time constant, voltage sag ratios, rheobase, resting membrane potential, input resistance, and SFA ratio. The results showed that the first principal component explained 92.28% of the variability in the data and the second component explained 7.14% of the variability. Despite largely different intrinsic physiology, the PCA analysis did not separate Y_1_R^+^ neurons with adapting or sustained firing patterns into non-overlapping clusters (**Figure 3H**). To probe this quantitatively, we used k-means cluster analysis. With K-means cluster analysis set to identify 2 clusters, we found that the majority of adapting neurons (69.70%) were part of cluster 1 and most sustained neurons (65%) fell into cluster 2 (**Figure 3I**). These data suggest that even though individual intrinsic properties differ between adapting and sustained neurons at the population level, these differences do not completely separate Y_1_R^+^ neurons into two different groups. This is likely due to the variability in intrinsic physiology within each neuron group, which often overlapped between sustained and adapting neurons. These results suggest that IC Y_1_R^+^ neurons encompass two or more classes of glutamatergic neurons and provide additional support for the observation that IC neurons are difficult to classify based on physiology alone (Reetz and Ehret, 1999; Peruzzi et al., 2000; Sivaramakrishnan and Oliver, 2001; Ono et al., 2005; Goyer et al., 2019; Kreeger et al., 2021).

### Y_1_R^+^ neurons form highly interconnected networks in the local IC

The IC is rich in local circuits, with anatomical reports indicating that most neurons have local axon collaterals (Oliver et al., 1991; Ito et al., 2016). However, very few studies have examined the functional organization of local circuits in the IC, and it is unknown how different populations of neurons contribute to IC local circuits. To investigate how Y_1_R^+^ neurons contribute to local circuits, we used viral transfections to express the excitatory opsin Chronos-GFP (Klapoetke et al., 2014) in Y_1_R^+^ neurons in one side of the IC, and then we targeted our recordings to either Y_1_R^+^ neurons or Y_1_R^-^ neurons in the transfected side of the IC. Since most glutamatergic IC neurons express Y_1_R and glutamatergic neurons represent ∼75 % of neurons in the IC (Oliver et al., 1994; Beebe et al., 2016), we estimate that there is ∼77 % chance that Y_1_R^-^ neurons are GABAergic neurons. (This estimate is based on previous reports that GABAergic neurons represent ∼25 % of IC neurons and our in situ hybridization result showing that tdTomato is expressed by 92.5 % of glutamatergic neurons in the IC: 25%/ (25% + 7.5%) = 77%). However, since no GABAergic marker was used to identify these cells, we will refer to these neurons as Y_1_R^-^ neurons.

First, we targeted recordings to Y_1_R^+^ neurons in acute brain slices. We found that activation of Y_1_R^+^ terminals with brief pulses of blue light (0.5 – 5 ms) elicited EPSPs in most Y_1_R^+^ neurons (**Figure 4A-C**; 14 out of 16 neurons recorded). Neurons in the IC can express NMDA receptors that are relatively insensitive to Mg^2+^ block and therefore, readily conduct current at resting membrane potential (Wu et al., 2004; Goyer et al., 2019; Drotos et al., 2023). To test whether such NMDA receptors are present at Y_1_R^+^ synapses onto Y_1_R^+^ neurons, we bath applied 10 μM of NBQX, which is an AMPA receptor antagonist. In the presence of 10 μM NBQX, the light-evoked EPSP was incompletely blocked in 10 out of 14 cells, suggesting the presence of a NMDA component (amplitude in control condition: 4.6 mV ± 3.0 mV; amplitude in NBQX: 0.6 mV ± 0.5 mV; LMM: β = -4.01, 95% CI [-5.33 -2.69], *p* = 1.36e-05, n = 14. **Figure 4B-C**). The EPSPs were completely abolished when 50 μM D-AP5, an NMDA receptor antagonist, was applied (LMM: β = -4.57, 95% CI [-6.21 -2.94], *p* = 1.88e-05, n = 4. **Figure 4B-C**).

**Figure 3.**
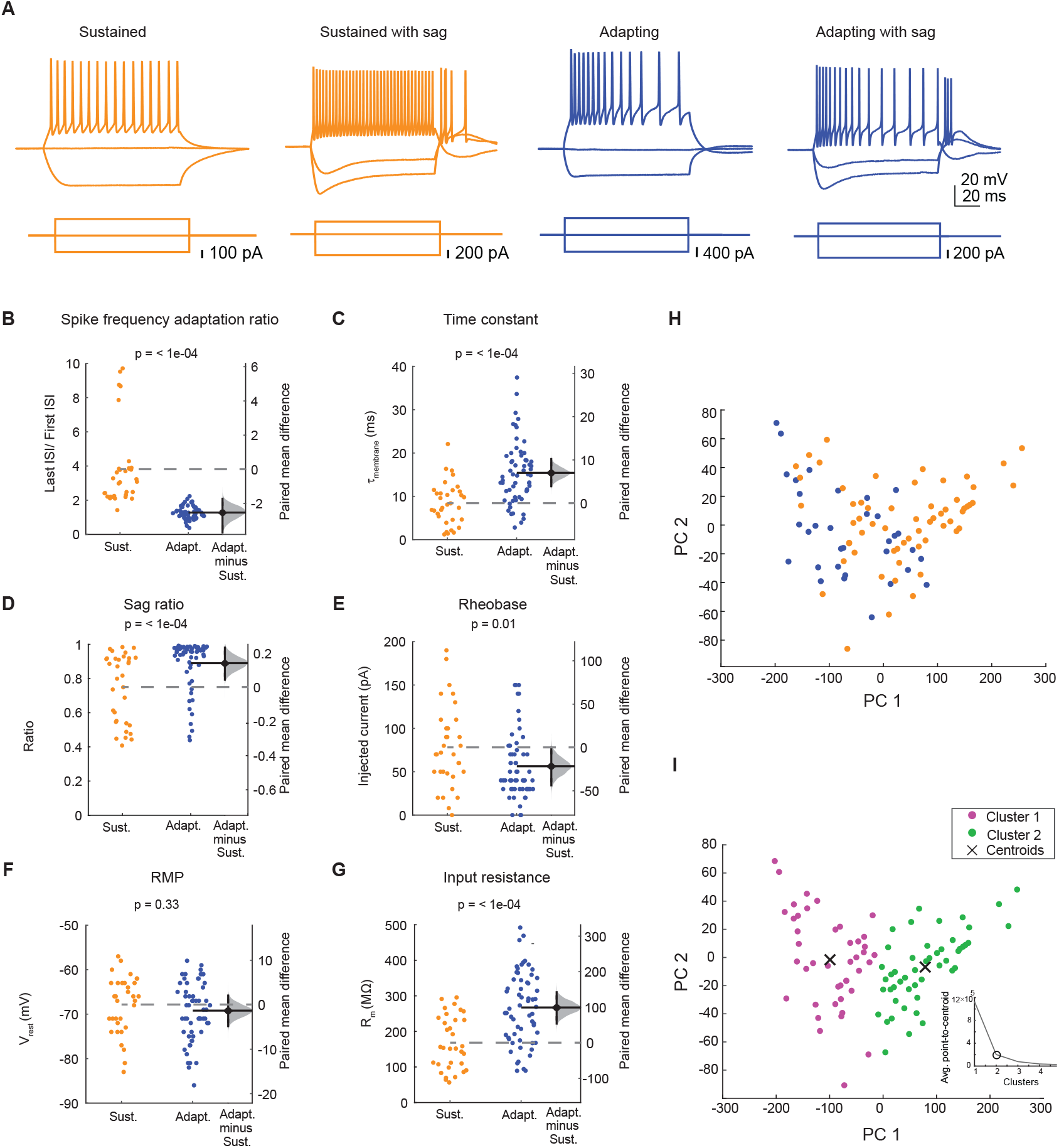
Y_1_R^+^ neurons exhibit sustained or adapting firing patterns. **A**. Y_1_R^+^ neurons exhibited different combinations of firing patterns and hyperpolarization-induced sag. Orange indicates neurons with a sustained firing pattern with or without voltage-dependent sag and blue indicates neurons with an adapting firing pattern with or without voltage-dependent sag. **B – G**. Sustained and adapting neurons exhibited heterogenous intrinsic physiological properties. **B**, Spike frequency adaptation ratio. **C**, Membrane time constant. **D**, Voltage-dependent sag ratio. **E**, Rheobase. **F**, Resting membrane potential. **G**, Input resistance. Dashed gray lines represent the level of zero difference in the mean difference plots. **H**. Principal components analysis showed that the distributions of adapting (blue) and sustained (orange) neurons overlapped. **I**. Separation of Y_1_R^+^ neurons into two clusters using k-means cluster analysis yielded clusters that did not match those predicted from sustained and adapting firing patterns (compare H and I). Magenta and green dots represent the two different clusters. 69.7% of adapting neurons fell into cluster 1 and 65.0 % of sustained neurons fell into cluster 2. The number of clusters used for analysis was defined using the elbow analysis, which showed that the addition of a third cluster did little to improve the separation between cluster centroids (insert graph).

**Figure 4.**
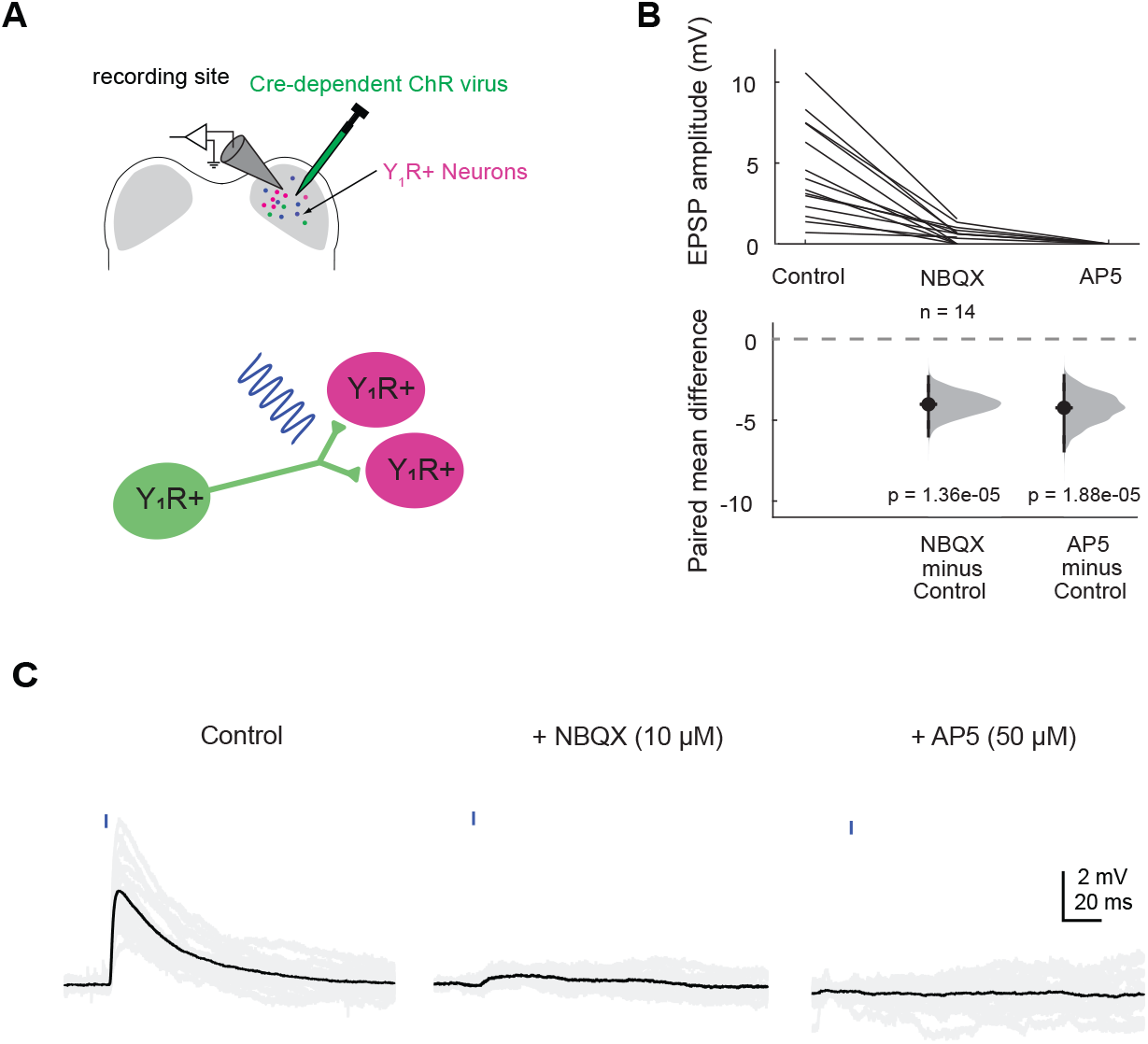
Y_1_R^+^ neurons synapse onto other Y_1_R^+^ neurons. **A**. Cartoon representing the experimental setup. A Cre-dependent AAV was injected into one side of the IC to drive Chronos expression in Y_1_R^+^ neurons. After allowing 2 – 4 weeks for opsin expression, recordings were targeted to Y_1_R^+^ neurons in the transfected side of the IC. A brief pulse of blue light was used to activate Y_1_R^+^ terminals. **B**. Light pulses elicited EPSPs of varying amplitudes. These EPSPs were blocked by 10 μM NBQX in 10 out of 14 cells tested. The remaining EPSPs were abolished after application of 50 μM AP5. The dashed gray line indicates the level of zero difference in the paired mean difference plot. **C**. Example traces of optogenetically evoked EPSPs recorded from a Y_1_R^+^ neuron in the IC ipsilateral to the injection site. Black traces represent average responses and gray traces represent individual sweeps.

Next, we targeted our recordings to Y_1_R^-^ neurons that were identified by lack of tdTomato expression. We found that presentation of 1 -2 ms light pulses elicited EPSPs in 6 out of 6 cells tested. Interestingly, after application of 10 μM NBQX, only one cell had an incompletely blocked EPSP, suggesting the presence of an NMDA component (**Figure 5A-C**, amplitude in control condition: 2.8 mV ± 1.0 mV; amplitude in NBQX: 0.1 mV ± 0.3 mV; LMM: β = -4.01, 95% CI [-3.51 -1.93], *p* = 5.23e-05, n = 6). Application of 50 μM D-AP5 completely abolished the EPSP. Because the NMDA component was present in only one cell, we did not run statistical analysis on these data, but we show the raw data in **Figure 5C**.

**Figure 5.**
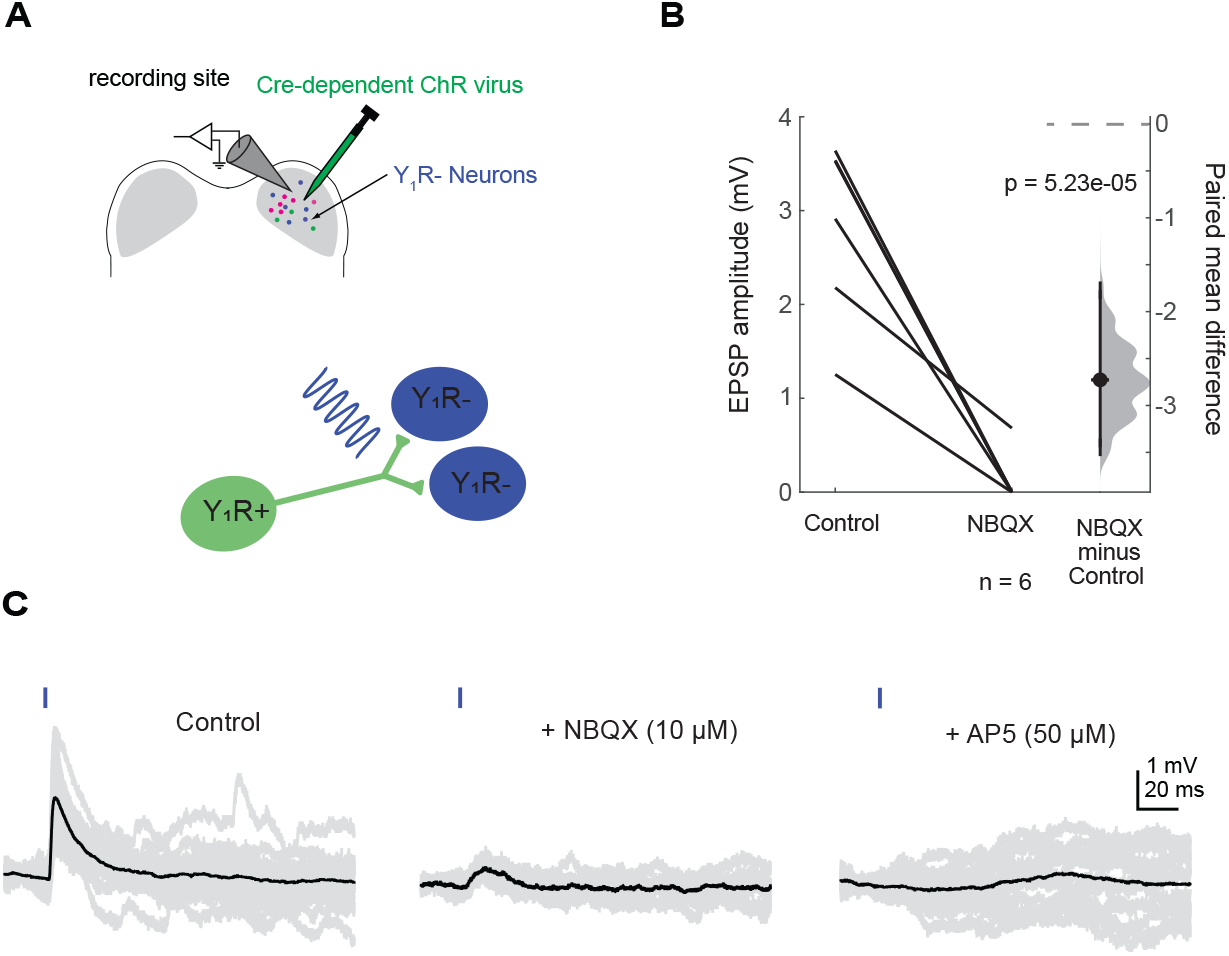
Y_1_R^+^ neurons synapse onto other Y_1_R^-^ neurons. **A**. Cartoon representing the experimental setup. A Cre-dependent AAV was injected into the right IC to drive Chronos expression in Y_1_R^+^ neurons. After allowing 2 – 4 weeks for virus expression, recordings were targeted to Y_1_R^-^ neurons in the transfected side of the IC. A brief pulse of blue light was used to activate Y_1_R^+^ terminals. **B**. Light pulses elicited EPSPs of varying amplitudes. Application of 10 μM NBQX abolished EPSPs in 5 out of 6 cells tested. In the sixth cell, the remaining EPSP was abolished with application of 50 μM AP5 (data not shown on graph). **C**. Example traces of optogenetically evoked EPSPs recorded from a Y_1_R^-^ neuron in the IC ipsilateral to the injection site. EPSPs are from the only Y_1_R^-^ neuron that exhibited an NMDA component in its response. Black traces represent average responses and gray traces represent individual sweeps.

### Y_1_Rs synapses have modest short-term synaptic plasticity

To investigate whether inputs mediated by Y_1_R^+^ neurons exhibit short-term synaptic plasticity, we recorded from 6 Y_1_R^+^ neurons and 6 Y_1_R^-^ neurons while presenting blue light pulses to activate Y_1_R^+^ inputs. For most recordings (8 out of 12 cells), we used viral transfections to drive the expression of a soma-targeted variant of the excitatory opsin ChroME in Y_1_R^+^ neurons (Mardinly et al., 2018). Soma-targeted ChroME allowed us to study synaptic events generated by eliciting action potentials in the cell body rather than direct depolarization of synaptic terminals, providing a more physiological condition for short-term synaptic plasticity experiments.

In four mice, we used viral transfections to express Chronos in Y_1_R^+^ neurons. In these neurons, vesicular release was likely achieved by direct depolarization of synaptic terminals. Since no difference was observed between these approaches, the data were combined.

We used 20 Hz train stimulation to elicit EPSPs (**Figure 6**) and obtained the PPR by dividing the average peak amplitude of the second, third or fourth EPSP over the average peak amplitude of the first EPSP (see Materials and Methods for details). Interestingly, in contrast to what has been reported in ascending projections from the lemniscal pathway (Wu et al., 2004), we saw little short-term synaptic plasticity when recording either from Y_1_R^+^ or Y_1_R^-^ neurons. The PPR values when recording from Y_1_R^+^ neurons were: EPSP_2_/EPSP_1_ = 0.95 ± 0.17, EPSP_3_/EPSP_1_ = 0.85 ± 0.14, EPSP_4_/EPSP_1_ = 0.95 ± 0.31 and EPSP_5_/EPSP_1_ = 0.86 ± 0.22 (mean ± SD). PPR values were not significantly different than 1 (*one-sample Student’s t test* comparing data to mean = 1: 1 *vs* EPSP_2_/EPSP_1_ *p* = 0.50; 1 *vs* EPSP_3_/EPSP_1_ *p* = 0.06; 1 *vs* EPSP_4_/EPSP_1_ *p* = 0.73; 1 *vs* EPSP_5_/EPSP*1 p* = 0.19; Bonferroni-corrected α = 0.0125). The PPR values determined from Y_1_R^-^ neurons were: EPSP_2_/EPSP_1_ = 1.11 ± 0.13, EPSP_3_/EPSP_1_ = 1.04 ± 0.16, EPSP_4_/EPSP_1_ = 1.00 ± 0.09 and EPSP_5_/EPSP_1_ = 0.93 ± 0.12. PPR values in Y_1_R^-^ neurons were not significantly different than 1 (*one-sample Student’s t test* comparing data to mean = 1: 1 *vs* EPSP_2_/EPSP_1_ p = 0.08; 1 *vs* EPSP_3_/EPSP_1_ *p* = 0.56; 1 *vs* EPSP_4_/EPSP_1_ *p* = 0.98; 1 *vs* EPSP_5_/EPSP_1_ *p* = 0.25. Bonferroni-corrected α = 0.0125). These data suggest that Y_1_R^+^ synapses in local IC circuits possess mechanisms that result in a good balance between short-term synaptic depression and short-term synaptic facilitation, thereby resulting in little short-term synaptic plasticity overall.

**Figure 6.**
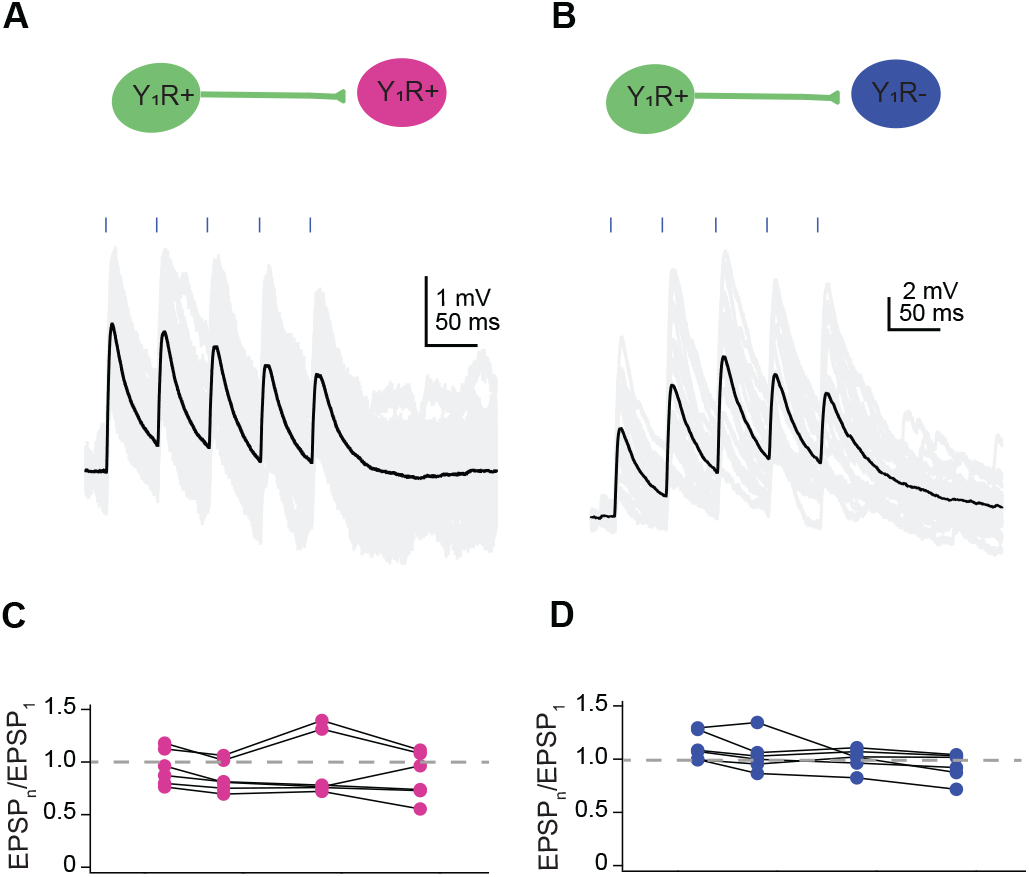
Y_1_R^+^ synapses exhibit moderate short-term synaptic plasticity. **A,B**. Example traces from a Y_1_R^+^ neuron (**A**) and a Y_1_R^-^ neuron (**B**) showing EPSPs evoked by 20 Hz trains of light pulses. Black traces represent average responses and gray traces represent individual sweeps. **C,D**. Plots of the paired pulse ratios (PPR) for Y_1_R^+^ synapses onto Y_1_R^+^ neurons (**C**) and Y_1_R^-^ neurons (**D**) reveal little short-term plasticity. Each dot represents (average EPSP_n_)/(average EPSP_1_) for an individual cell. Dashed gray lines indicate PPR of one.

### NPY decreases recurrent excitation in the IC

NPY is one of the most abundant neuropeptides in the brain and has been shown to modulate neuronal circuits in many brain regions (Colmers and Bleakman, 1994; Gutman et al., 2008; van den Pol et al., 2009; van den Pol, 2012). In the hippocampus, for example, NPY decreases recurrent excitation (Tu et al., 2005). Our data show that most glutamatergic neurons in the IC express Y_1_R and that Y_1_R^+^ neurons are highly interconnected with other Y_1_R^+^ and Y_1_R^-^ neurons in local IC circuits. This raises the hypothesis that NPY signaling regulates the excitability of local circuits in the IC. To test this hypothesis, we used a combination of pharmacology and optogenetics to generate recurrent excitation in vitro. For that, Y_1_R-Cre x Ai14 mice were injected with viruses to selectively express soma-targeted ChroME (10 out of 11 cells) or Chronos (1 out of 11 cells) in Y_1_R^+^ neurons. To increase the probability of recurrent excitation during slice recordings, we included 5 μM gabazine (GABA_A_ receptor antagonist) and 1 μM strychnine (glycinergic receptor antagonist) in the ACSF (Tu et al., 2005). When Y_1_R^+^ neurons were activated with a brief flash of blue light, prolonged trains of excitatory events, consistent with recurrent excitation, were observed in both Y_1_R^+^ and Y_1_R^-^ neurons (**Figure 7**).

**Figure 7.**
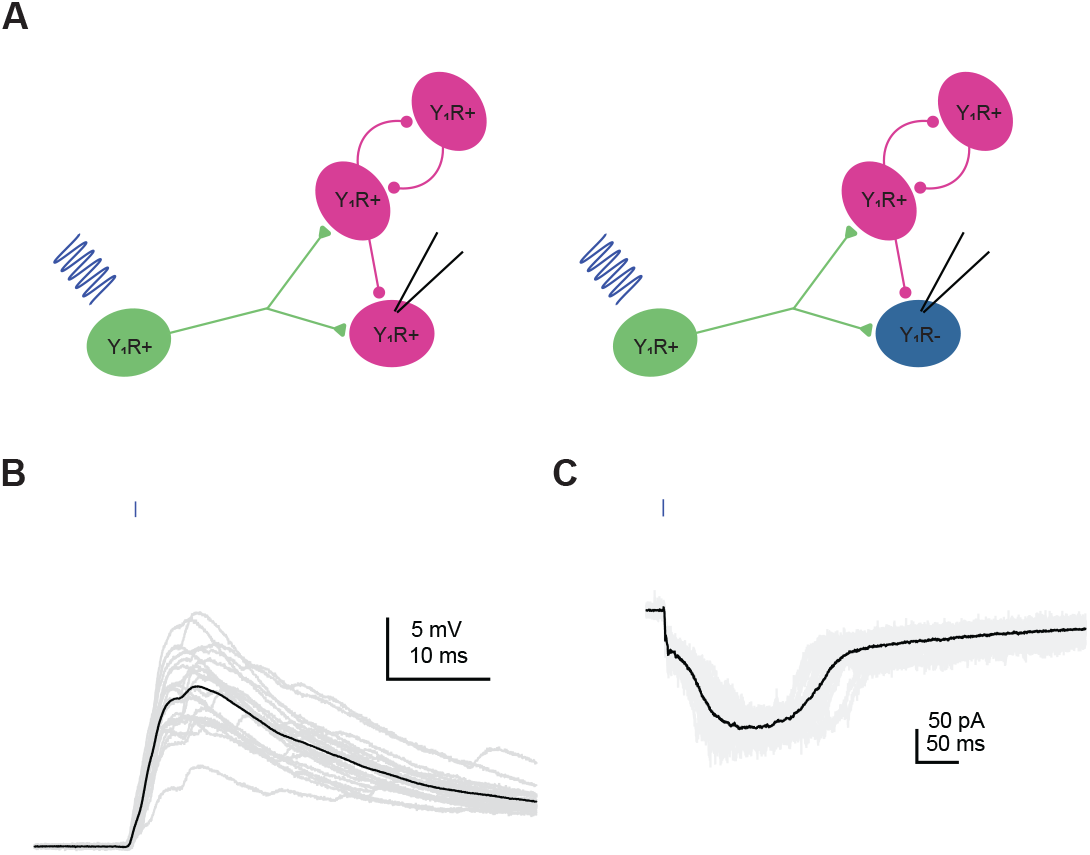
Activation of Y_1_R^+^ neurons elicits recurrent excitation in the IC. **A**. Cartoon showing experimental design. Viruses expressing Chronos or ChroME were used to transfect Y_1_R^+^ neurons. In the presence of inhibitory synaptic blockers (5 μM gabazine and 1 μM strychnine), optogenetic activation of Y_1_R^+^ neurons elicited prolonged periods of recurrent excitation. Recordings were targeted both to Y_1_R^+^ and Y_1_R^-^ neurons. **B,C**. Examples of recurrent excitation from a current-clamp recording (**B**) and a voltage clamp recording (**C**). Example trace in B is from a Y_1_R^+^ neuron and example trace in C is from a Y_1_R^-^ neuron.

**Figure 8.**
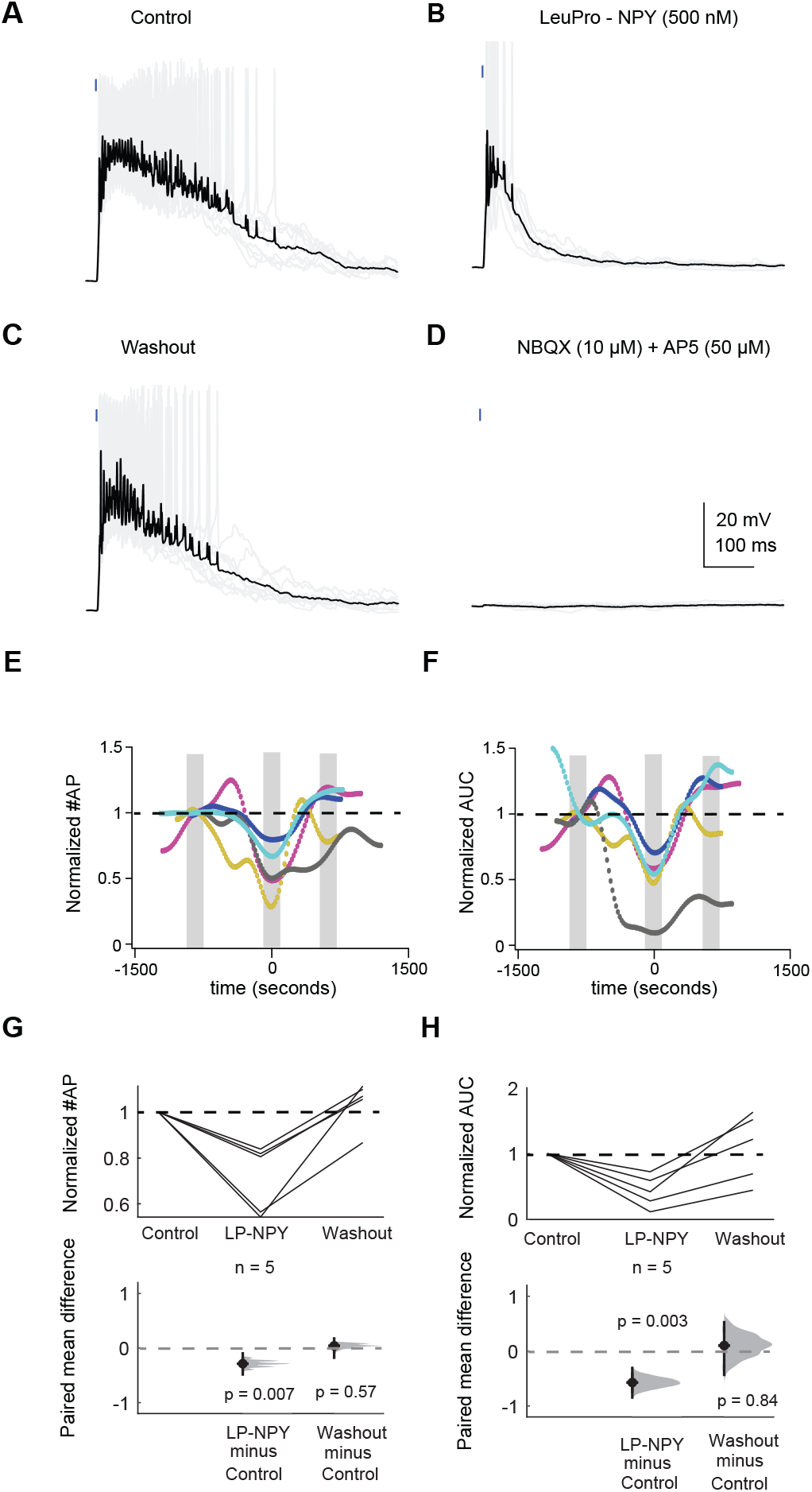
Application of LP-NPY decreases action potentials elicited by recurrent excitation. **A-D**. In a recording from a Y_1_R^+^ neuron, activation of other Y_1_R^+^ neurons using a brief light pulse elicited recurrent excitation that resulted in action potentials (**A**). Bath application of the Y_1_R agonist, LP-NPY (500 nM), decreased recurrent excitation resulting in a decrease in action potential number (**B**). This effect was reversed during washout (**C**). Recurrent excitation was completely abolished by application of 10 μM NBQX and 50 μM D-AP5 (**D**). **E**. Graph shows normalized action potential number across the total duration of the recordings. Data were normalized to the most negative point during LP-NPY application. Grey bars indicate the data points that were analyzed for baseline, LP-NPY application and washout. Black dashed line represents the level of one. Each color represents one individual cell, and each dot represents a single trial. Trails were run at 20 second intervals. **F**. Graph showing normalized area under the curve across the recorded cells. Colors correspond to cells in E. Data were normalized to the most negative point during LP-NPY application. Grey bars represent the data points that were analyzed for baseline, LP-NPY application and washout. Black dashed line represents the level of one. **G,H**. Application of LP-NPY decreased the number of action potentials (**G**) and the area under the curve (**H**) observed in response to light pulses, indicating that LP-NPY inhibited recurrent excitation. Black dashed lines represent the level of one, and dashed gray lines represent the level of zero difference for the paired mean difference plots.

First, we targeted whole-cell current clamp recordings to 2 Y_1_R^+^ neurons and 3 Y_1_R^-^ neurons. A brief presentation of blue light (0.5 – 5 ms) elicited recurrent excitation that elicited trains of action potentials in the recorded neurons (**Figure 8A**). Bath application of 500 nM LP-NPY, a Y_1_R agonist, decreased recurrent excitation, decreasing the number of action potentials elicited by the blue light pulse (LMM: β = -0.43, 95% CI [-0.66 -0.19, *p* = 0.007, n = 5. **Figure 8B,E,G**). After a 10 – 20 min washout in control ACSF, the number of action potentials elicited by recurrent excitation was similar to control (LMM: β = -0.07, 95% CI [-0.30 0.16, *p* = 0.57, n = 5, **Figure 8C,E,G**). In 4 out of 5 cells we applied NBQX and AP5 to verify that the recurrent excitation was synaptically driven (**Figure 8D**). In only one cell (a Y_1_R^+^ neuron) a brief EPSP persisted in the presence of synaptic blockers (amplitude: 13.5 ± 0.3 mV, half-width: 13.8 ± 0.6 ms; data not shown). However, since it was a brief EPSP it could not explain the long-lasting recurrent excitation observed in this experiment. We previously showed that NPY can directly hyperpolarize the membrane potential of Y_1_R^+^ neurons (Silveira et al., 2020), however, here we did not see a correlation between resting membrane potential and the number of action potential elicited by activation of Y_1_R^+^ neurons (data not shown, linear correlation test; cell #1 Y_1_R^+^ : r2 = -0.071, *p* = 0.004, cell #2 Y_1_R^+^: r2 = 0.085, *p* = 0.005, cell #3 Y_1_R^-^: r2 = -0.001, *p* = 0.690, cell #4 Y_1_R^-^: r2 = -0.018, *p* = 0.181, cell #5 Y_1_R^-^: r2 = -0.074, *p* = 0.005).

We next calculated the area under the curve. As detailed in the Methods, we used a median-filter to isolate the steady depolarization underlying the train of action potentials before determining the area under the curve. Application of LP-NPY led to a decrease in the area under the curve (LMM: β = -0.44, 95% CI [-0.65 -0.23, *p* = 0.003, n = 5) and the effect was reversed during washout (LMM: β = -0.005, 95% CI [-0.21 0.20, *p* = 0.84, n = 5. **Figure 8F,H**). Graphs in **Figure 8E,F** represent the changes in action potential number and area under the curve across the duration of each recording.

Next, we performed voltage-clamp experiments to investigate how NPY signaling shapes recurrent excitatory currents in the local IC. In recordings from 4 Y_1_R^+^ neurons and 2 Y_1_R^-^ neurons, we used optogenetics to activate Y_1_R^+^ neurons and saw long-lasting excitatory currents in the recorded neurons (**Figure 9A-C**). Application of LP-NPY decreased the recurrent excitatory current (LMM: β = -0.25, 95% CI [-0.45 -0.06, *p* = 0.03, n = 6. **Figure 9E,F**). Washout responses were obtained in 4 cells, showing a reversal of the LP-NPY effect (LMM: β = 0.14, 95% CI [-0.08 0.37, *p* = 0.25, n = 4). In these 4 cells, we next applied NBQX (10 μM) + AP5 (50 μM) to verify that excitatory responses were synaptically driven (**Figure 9D**). The response was completely abolished in 3 out of 4 cells tested (data not shown). In the fourth cell, the half-width of the remaining event was very brief (0.58 ± 0.01 ms), and therefore it could not explain the prolonged excitatory currents observed in that cell. In one additional cell, the recurrent excitation increased throughout the recording session and therefore it was excluded from the final analysis. Together, these data suggest that NPY signaling plays an important role in regulating excitatory networks in local IC circuits.

**Figure 9.**
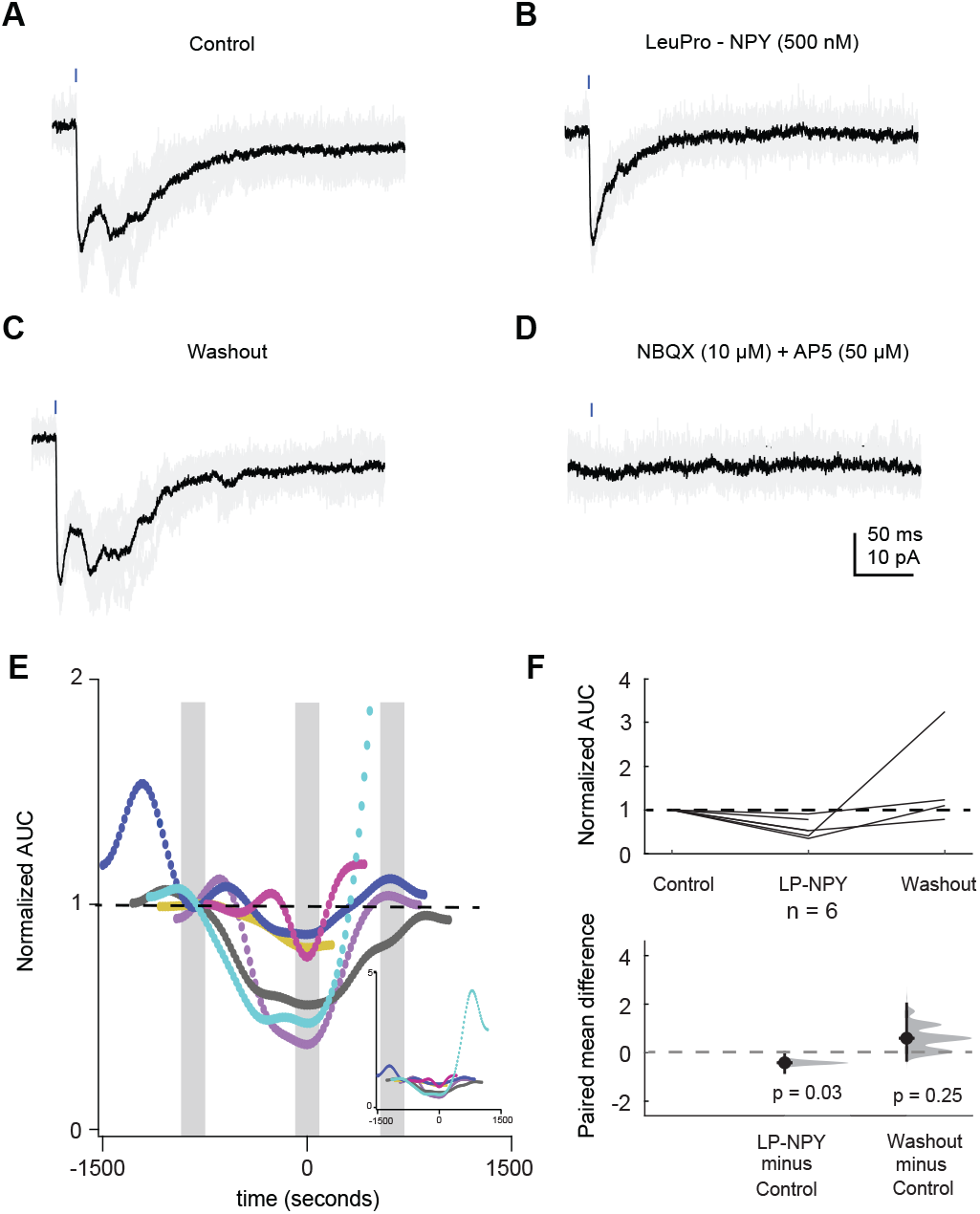
Application of LP-NPY decreases recurrent excitatory current. **A-D**. Activation of Y_1_R^+^ neurons by a brief light pulse elicited recurrent excitatory currents in an example recording from a Y_1_R^+^ neuron (**A**). Bath application of LP-NPY (500 nM) decreased recurrent excitation (**B**). This effect was reversed during washout (**C**). The recurrent excitatory current was abolished by 10 μM NBQX and 50 μM D-AP5 (**D**). **E**. Graph showing normalized area under the curve across 6 cells. Each dot represents one trial, with trial responses collected at 20 second intervals. Data were normalized to the most negative point during LP-NPY application. Grey bars indicate the data points that were analyzed for baseline, LP-NPY application and washout. Black dashed line represents the level of one. Each color represents one individual. The graph was truncated along the y-axis to better show the changes observed, removing some of the later responses for the cell in cyan, which ran-up during the washout period. The insert graph shows the full y-axis. **E**. Graph of normalized data showing that application LP-NPY (500 nM) decreased the area under the curve. Black dashed line represents the level of one, and the dashed gray line represents the level of zero difference for the paired mean difference plot.

## Discussion

Here, we showed that nearly 80% of glutamatergic neurons in the IC express mRNA encoding the *NPY Y*_*1*_*R*. Using targeted recordings and optogenetics, we found that Y_1_R^+^ neurons provide excitatory input to most other Y_1_R^+^ and Y_1_R^-^ neurons in the IC and therefore form highly interconnected networks within local IC circuits. Excitatory synapses between Y_1_R^+^ neurons and other IC neurons exhibited modest short-term synaptic plasticity, suggesting a balance between synaptic facilitation and depression that results in a stable strength of excitatory synaptic signaling in local circuits. Furthermore, we found that NPY shaped local excitation in the IC by inhibiting recurrent excitatory circuits. Thus, our data provide functional evidence that IC glutamatergic neurons form densely interconnected local circuits and indicate that NPY is a major modulator of local circuit operations in the auditory midbrain.

### Most excitatory neurons in the IC express the NPY Y_1_R

Neurons that express the Y_1_R are widely distributed in the brain (Eva et al., 2006). Y_1_Rs are mainly postsynaptic (Wahlestedt et al., 1986; Kopp et al., 2002; Fu et al., 2004), but can also be found at presynaptic sites (Dumont et al., 1998; Glass et al., 2002). The neurotransmitter content of Y_1_R^+^ neurons vary across brain regions. In the amygdala and hypothalamus, Y_1_R can be expressed by GABAergic and glutamatergic neurons (Roseberry et al., 2004; Rostkowski et al., 2009; Wittmann et al., 2013). In the prefrontal cortex (Vollmer et al., 2016) and spinal cord (Nelson et al., 2019), most Y_1_R^+^ neurons are glutamatergic. Strikingly, in the IC, which is well known for its neuronal diversity (Peruzzi et al., 2000; Palmer et al., 2013; Beebe et al., 2016), *Y*_*1*_*R*^*+*^ neurons almost completely overlapped with *Vglut*^*2+*^ neurons (**Figure 1**) and very rarely co-labeled with GAD67 (Silveira et al., 2020). The fact that NPY is expressed in approximately one-third of GABAergic IC neurons (Silveira et al., 2020) while Y_1_R is expressed in most glutamatergic IC neurons represents a major step towards understanding the functional organization of neuronal circuits in the IC. Since our results indicate that Y_1_R^+^ neurons encompass multiple classes of IC glutamatergic neurons, it will be important to determine whether activation of Y_1_Rs by NPY signaling inhibits different classes of IC excitatory neurons in subtly different ways or whether NPY signaling broadly and equally dampens all local excitatory activity in the IC.

### Y_1_R^+^ neurons form highly interconnected local circuits in the IC

Anatomical studies suggest that most IC neurons have local axon collaterals (Oliver et al., 1991; Saldana and Merchan, 2005; Chen et al., 2018). Previous studies using laser scanning glutamate uncaging provided functional evidence of local excitatory and inhibitory networks in the IC of young mice (P2 – P22) (Sturm et al., 2014, 2017). Additionally, a recent study showed that descending input from auditory cortex to the shell IC elicits net inhibitory responses by driving local glutamatergic neurons to activate local networks of GABAergic neurons that in turn synapse broadly onto other shell IC neurons (Oberle et al., 2022). Here most of our recordings were targeted to central nucleus of the IC and we found that intrinsic Y_1_R^+^ projections are very common, since only 2 out of 45 recorded cells did not receive local inputs from Y_1_R^+^ neurons. This suggests that Y_1_R^+^ neurons, which represent most of the glutamatergic IC cells, form much more highly interconnected networks in the IC than previously expected.

Glutamatergic synapses in the IC often activate both AMPA and NMDA receptors in postsynaptic IC neurons (Ma et al., 2002; Wu et al., 2004; Goyer et al., 2019). Here we showed that 10 out of 14 Y_1_R^+^ neurons exhibited an NMDA receptor component in their EPSPs at resting membrane potential. This could indicate that local synapses are located on distal dendrites or dendritic spines where activation of AMPA receptors might be sufficient to remove voltage-dependent Mg^2+^ block of NMDA receptors. However, a recent study from our lab showed that many IC neurons express GluN2C and/or GluN2D NMDA receptor subunits (Drotos et al., 2023), which are relatively insensitive to Mg^2+^ block (Siegler Retchless et al., 2012). Here, recordings that were targeted to Y_1_R^-^ cells, which are likely GABAergic, rarely exhibited an NMDA receptor component (1 out of 6 cells). The different NMDA receptor expression between Y_1_R^+^ and Y_1_R^-^ neurons may influence auditory computations in the IC by differently affecting synaptic integration in glutamatergic versus GABAergic neurons (Drotos et al., 2023).

Most synapses undergo dynamic changes during successive release events that result in a temporary increase or decrease in synaptic strength known as short-term synaptic depression or short-term synaptic facilitation. Short-term synaptic plasticity changes the efficacy of a synapse and directly influences neuronal computations, affecting for example, neuronal gain (Dittman et al., 2000; Rothman et al., 2009; Barri et al., 2022). Short-term synaptic plasticity has been widely studied in the central auditory pathway (Friauf et al., 2015; Romero and Trussell, 2021), however very few studies have been done in the IC (Wu et al., 2004; Kitagawa and Sakaba, 2019). Because axons from multiple sources overlap in the IC, it is challenging to use electrical stimulation to activate single sources of input to IC neurons and even more challenging to separate local from external sources of input. Here we used optogenetics to selectively stimulate Y_1_R^+^ neurons in the local IC. We showed that, in contrast to ascending inputs (Wu et al., 2004), glutamatergic synapses in the local IC exhibited modest short-term synaptic plasticity. This suggests that synaptic mechanisms in the local IC favor stable synaptic strength at excitatory synapses. Short-term synaptic plasticity has been hypothesized to contribute to temporal selectivity influencing, for example, sensory adaptation in which the response to a second auditory stimulus is decreased compared to the first stimulus (Phillips et al., 1989; Natan et al., 2017; Motanis et al., 2018; Seay et al., 2020; Valdés-Baizabal et al., 2021). The difference between short-term synaptic plasticity between ascending inputs and local synapses may indicate a shift in the influence from ascending to local synapses during sustained stimuli.

### NPY signaling dampens recurrent excitation in the IC

The IC receives several neuromodulatory inputs that mostly come from non-auditory sources outside the IC (Hurley and Pollak, 1999; Motts and Schofield, 2009; Rivera-Perez et al., 2021; Hoyt et al., 2019). Because NPY neurons are located within the IC, NPY signaling is likely to be much more heavily influenced by the ascending auditory pathway than other neuromodulatory inputs to the IC. We previously showed that NPY signaling dampens the excitability of Y_1_R expressing neurons (Silveira et al., 2020). Our current results extend this finding by showing that, since most glutamatergic IC neurons express Y_1_R, NPY is positioned to be a major regulator of excitatory neuronal circuits in the IC.

The importance of NPY signaling has been shown in many brain regions. For example, in the hippocampus, NPY is known as an endogenous anti-epileptic agent that decreases excitability during seizures (Colmers and El Bahh, 2003; Giesbrecht et al., 2010). Interestingly, the role of Y_1_Rs in modulating seizures in the hippocampus is debated. Some studies suggest that Y_1_Rs play a permissive role in seizures (Vezzani et al., 1999) and that the anti-epileptic role of NPY is likely to be mediated by presynaptic Y2Rs (Colmers et al., 1988; Colmers and El Bahh, 2003; Vezzani and Sperk, 2004). Here, we showed that activation of Y_1_Rs decreases recurrent excitation in the IC. This contrast could be explained by the fact that, in the hippocampus, Y_1_Rs can be auto-receptors expressed by NPY neurons (Paredes et al., 2003). In the IC, Y_1_Rs are expressed in glutamatergic neurons and are not expressed by NPY neurons, which are GABAergic (Silveira et al., 2020). Given that the IC is prone to hyperexcitability (Wei, 2013; Xiong et al., 2017) and that NPY expression in the IC is increased in a rat model of audiogenic seizure (Damasceno et al., 2020), NPY may play a major role in shaping excitatory/inhibitory balance in the IC.

Noise induced hearing loss can lead to immediate and long-term changes in the activity of IC neurons (Wang et al., 1996; Dong et al., 2010). Noise-induced plasticity in the IC is hypothesized to be a compensatory response to decreased excitatory drive from the periphery (Chambers et al., 2016). This phenomenon is known as enhanced central gain and, when the enhancement goes too far, it can produce maladaptive plasticity that results in clinical conditions such as tinnitus or hyperacusis (Berger and Coomber, 2015; McGill et al., 2022). Given that NPY strongly decreases excitability in the IC, it is not unreasonable to think that NPY signaling may limit enhanced gain in the IC after hearing loss. In fact, a recent study showed that NPY expression is increased in lateral olivocochlear neurons after noise exposure (Frank et al., 2023). In future studies, we plan to directly assess how NPY signaling contributes to neural plasticity in the IC following noise-induced hearing loss.

Together, our data suggest that NPY signaling is a critical modulator of local circuits in the IC, with potential impacts on auditory computations, central gain control, and plasticity following hearing loss.

## Acknowledgements

We thank Bo Duan, Susan Shore and Pierre Apostolides for helpful discussions and advice. We also thank Yoani Herrera for help with cartoon illustrations. This work was supported by National Institutes of Health Grants K99 DC019415 (MAS) and R01 DC018284 (MTR), National Science Foundation Graduate Research Fellowship Program Grant DGE 1256260 (ACD), and National Science Foundation Grant 2243919 (TMP).

## Author Contributions

MAS and MTR designed the research. MAS, ACD, TMP, TSV and AB performed experiments. MAS and MTR analyzed data. MAS wrote the initial draft of the manuscript. MAS and MTR revised the manuscript. All authors approved the final version.

